# Cell cycle regulation of the psoriasis associated gene *CCHCR1* by transcription factor E2F1

**DOI:** 10.1101/2023.04.08.535951

**Authors:** Yick Hin Ling, Yingying Chen, Kwok Nam Leung, King Ming Chan, Wing Keung Liu

## Abstract

The coiled-coil alpha-helical rod protein 1 (CCHCR1) was first identified as a candidate gene in psoriasis and has lately been associated with COVID-19 susceptibility. Located within P-bodies and centrosomes, its exact cellular role and transcriptional control remain largely unknown. Here, we showed that *CCHCR1* shares a bidirectional promoter with its neighboring gene, *TCF19*. This bidirectional promoter is activated by the G1/S-regulatory transcription factor E2F1, and both genes are co-induced during the G1/S transition of the cell cycle. A luciferase reporter assay suggests that the short intergenic sequence, only 287 bp in length, is sufficient for the G1/S induction of both genes, but the expression of *CCHCR1* is further enhanced by the presence of exon 1 from both *TCF19* and *CCHCR1*. This research uncovers the transcriptional regulation of the *CCHCR1* gene, offering new perspectives on its function. These findings contribute to the broader understanding of diseases associated with CCHCR1 and may serve as a foundational step for future research in these vital medical fields.

## Introduction

*CCHCR1* (coiled-coil alpha-helical rod protein 1) is a candidate gene for psoriasis, a skin condition affecting 1% to 2% of the population [1]. CCHCR1 has been implicated in various cellular processes including keratinocyte proliferation and differentiation [2, 3], steroidogenesis [4, 5], myogenic differentiation [6], and cytoskeleton organization [7]. Studies from our lab have uncovered CCHCR1 as a protein associated with P-bodies and centrosome [8], suggesting its potential dual roles in regulating RNA metabolism and microtubules organization. This finding is further supported by RNA sequencing that highlights the role of CCHCR1 in related pathways [9], and knockdown of CCHCR1 caused centriole duplication defects and multipolar spindle formation [10]. Notably, the *CCHCR1* gene has recently been linked to alopecia areata [11], type-2 diabetes [12], and COVID-19 susceptibility [13], emphasizing the importance of understanding its cellular function and transcription regulation.

Psoriatic skin is characterized by hyper-proliferation and abnormal differentiation of the keratinocytes [14]. Early studies attempted to establish a correlation between the expression of CCHCR1 and cell proliferation, but the relationship is complex. For example, insulin and estrogen, which promote the growth of keratinocytes, up-regulate *CCHCR1* gene expression [5]; the immunosuppressant cyclosporine A inhibits the proliferation of keratinocytes and down-regulates *CCHCR1* [5]. Inactivation of the Rb pathway in neuronal cells results in cell cycle re-entry and up-regulation of *CCHCR1* [15]. On the contrary, serum starvation followed by re-feeding of cultured immortalized human HaCaT keratinocytes down-regulates *CCHCR1* [16]. The topoisomerase inhibitor camptothecin (CPT), which causes DNA damage, growth arrest, and cell apoptosis [17-20], up-regulates *CCHCR1* [21]. Notably, overexpression of CCHCR1 reduces keratinocyte proliferation [22]. Given that many of these experimental conditions affect both cell cycle progression and cell proliferation, it is imperative to explore the effect of the cell cycle on the transcriptional regulation of *CCHCR1*. In this study, we demonstrated that CCHCR1 shares a bidirectional promoter with its neighboring gene, *TCF19* (transcription factor 19). The activity of this bidirectional promoter is controlled by E2F1, a transcription factor involved in the G1/S cell cycle transition. In line with this, both *CCHCR1* and *TCF19* are co-induced during the G1/S transition. Our findings underscore the importance of understanding the cell cycle status of the cell for CCHCR1-associated diseases.

### E2F1 regulates *CCHCR1*-*TCF19* bidirectional promoter

The *CCHCR1* gene, located on human chromosome 6p21.3, is oriented head-to-head with its neighboring gene, *TCF19* (Fig 1A and B). We hypothesized that *CCHCR1* and *TCF19* share a bidirectional promoter since their transcriptional start sites are only 287 bp apart (Fig 1B). Typically, gene pairs with a bidirectional promoter are separated by less than 1000 bp [23]. The nucleotide sequence preceding the start codon of each gene reveals typical characteristics of a classical bidirectional promoter [23]. The region is TATA-less and contains a CpG island that overlaps the first exon of each gene (Fig 1B). Multiple putative transcription factor binding sites that are over-represented in the bidirectional promoter [24], such as GABP, MYC, E2F1, E2F4, NFY, and YY1 are found in the intergenic region of *CCHCR1*-*TCF19* (S1 Table and S2 Table). Notably, this region is highly enriched with E2F transcription factor binding motifs (S2 Table), a family of transcription factors that are instrumental in controlling the expression of genes that play a role in the G1/S transition and DNA synthesis [25]. Analysis of the microarray database [26] revealed a positive correlation between *CCHCR1* expression and genes associated with the G1/S transition or S phase of the cell cycle, including *E2F1* and *TCF19* (S3 Table). Furthermore, a positive correlation between *CCHCR1* and *TCF19* expression across various cell types suggests potential co-regulation (S3 Table).

**Fig 1.**
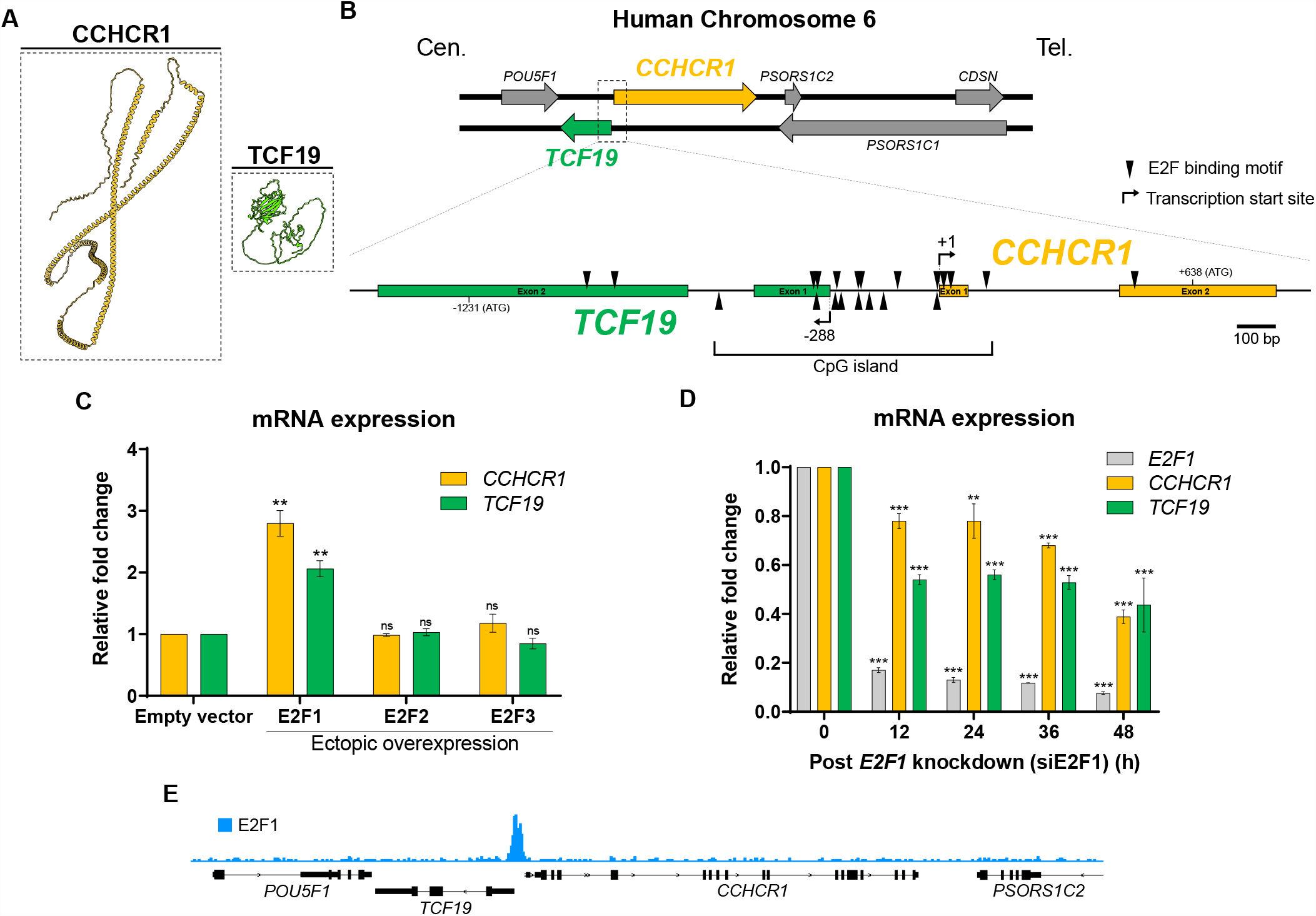
E2F1 induces the expression of *CCHCR1* and *TCF19*. (A) AlphaFold predicted structure of CCHCR1 and TCF19 proteins. (B) Genome arrangement of the *CCHCR1* and *TCF19* on chromosome 6. Cen.: Centromeric side, Tel.: Telomeric side. Putative E2F binding motifs (arrowhead) and CpG island shown. Numbers refer to the position related to the transcription start site (+1) of *CCHCR1*. (C) qPCR for the expression of *CCHCR1* and *TCF19* in HeLa cells transfected with E2F activators (E2F1, E2F2 or E2F3). (D) Down-regulation of *CCHCR1* and *TCF19* mRNA by E2F1 mRNA knockdown. (E) E2F1 shows high occupancy in the intergenic region of *CCHCR1*-*TCF19*. ChIP-seq data retrieved from ChIP-Atlas [SRX150563] [27].

The E2F transcription factor family consists of 8 members: E2F1-3 are activators, and E2F4-8 are repressors. To address the role of E2F in the expression of *CCHCR1* and *TCF19*, we measured the mRNA levels of *CCHCR1* and *TCF19* after ectopic overexpression of E2F1, 2, or 3 in HeLa cells. Among these three E2F activators, only the overexpression of E2F1 induced *CCHCR1* and *TCF19* expression (Fig 1C). In addition, the knockdown of *E2F1* mRNA by siRNA led to significant down-regulation of *CCHCR1* and *TCF19* (Fig 1D). Notably, E2F1 shows high occupancy in the *CCHCR1*-*TCF19* intergenic region (Fig 1E), as indicated by chromatin immunoprecipitation (ChIP) data in HeLa cells from ChIP-Atlas [27]. Therefore, E2F1 is functionally linked to the expression of *CCHCR1* and *TCF19*.

### *CCHCR1* and *TCF19* are up-regulated in the G1/S transition

Since E2F1 regulates genes in the G1/S transition [25], we explored the cell cycle expression of *CCHCR1* and *TCF19*. HeLa cells were synchronized at the G1/S transition by double thymidine block, and then released to allow cell cycle progression (Fig 2A and B). The expression patterns of *CCHCR1* and *TCF19* were closely correlated (Fig 2C); Their expression peaks at the G1/S transition, remains high during the S phase, and subsequently decreases during the transition to the G2/M phase. The expression levels continue to remain relatively low into the G1 phase (Fig 2C). Although cells begin to lose synchrony in subsequent rounds of the cell cycle, particularly from 16 h to 20 h after release (Fig 2A and B), we still observed a coordinated increase in the expression of *CCHCR1* and *TCF19* when cells approach the S phase (Fig 2C).

**Fig 2.**
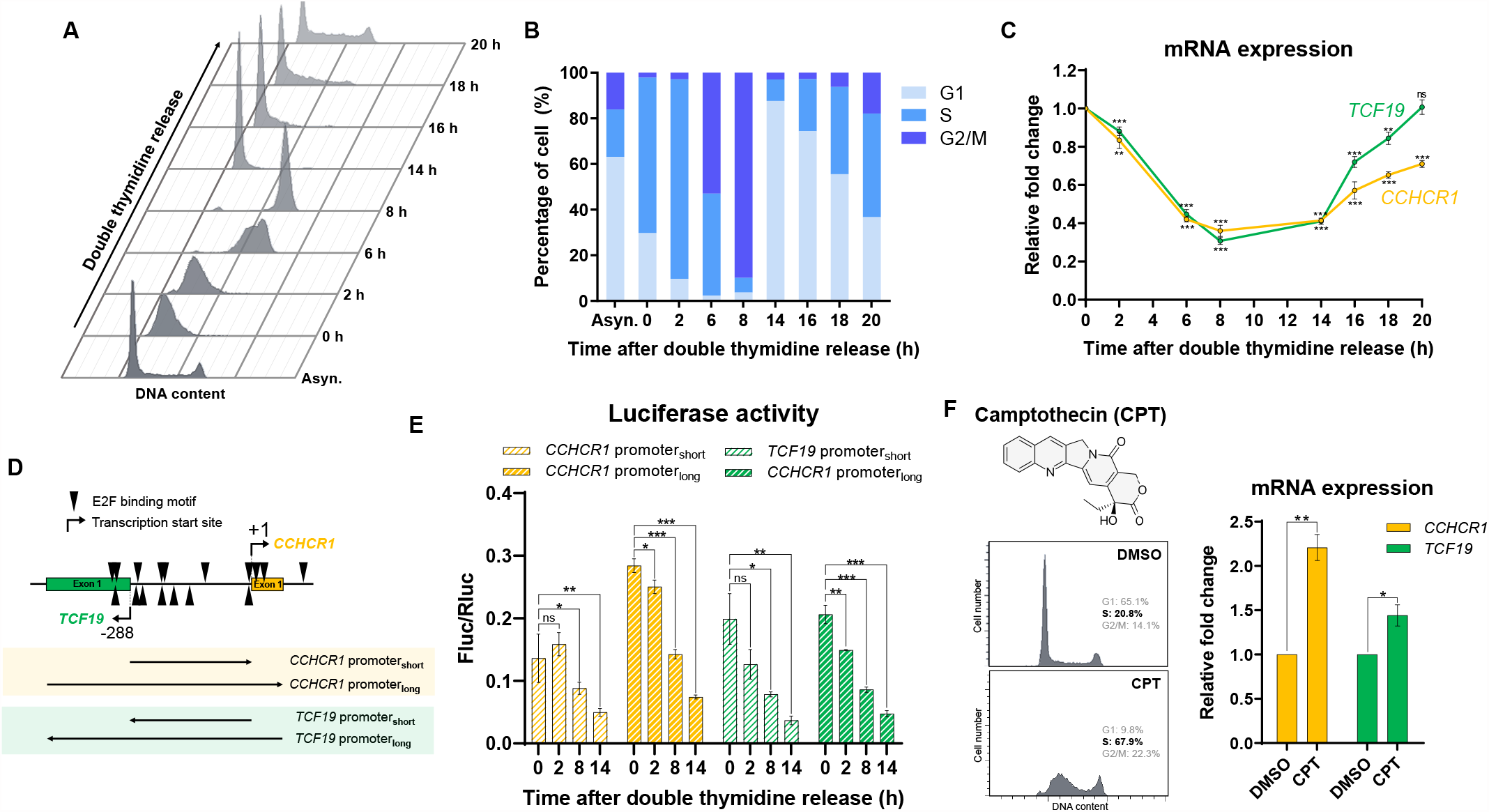
*CCHCR1* and *TCF19* are up-regulated in the G1/S transition. (A) DNA content and (B) percentage of cells in different cell cycle phases synchronized by double thymidine block and release. Asyn.: Asynchronous. (C) Relative expression of *CCHCR1* and *TCF19* in different cell cycle phases. (D) Promoter fragments of *CCHCR1*-*TCF19* bidirectional promoter for the dual luciferase assay. (E) Normalized luciferase activity (Fluc/Rluc) for *CCHCR1*-*TCF19* bidirectional promoter fragments. (F) DNA content (right) and relative expression of *CCHCR1* and *TCF19* (left) after 2 μM CPT treatment for 24 h.

To verify the activity of the *CCHCR1*-*TCF19* bidirectional promoter at the G1/S transition, we cloned promoter fragments into firefly luciferase (Fluc) reporter constructs and performed a dual luciferase assay to monitor their activity throughout the cell cycle. The short promoter fragments (287 bp) contain the intergenic sequence in either forward or reverse orientations (Fig 2D). The long promoter fragments (556 bp) additionally include exon 1 of both genes (Fig. 2D). These promoter fragments revealed high activities during the G1/S transition and S phase (Fig 2E). Specifically, the long fragment in the forward orientation for *CCHCR1* demonstrated higher activity across all cell cycle phases, unlike the corresponding fragment in the reverse orientation. Overall, we demonstrated that the activity of the *CCHCR1*-*TCF19* bidirectional promoter is up-regulated in the G1/S transition and maintained at a high level in S phase.

Recognizing the cell cycle regulation of *CCHCR1* and *TCF19* prompts us to further examine the effect of camptothecin (CPT) on their expression. CPT, a topoisomerase I inhibitor that induces cell cycle arrest at the G2 and S phases [17-20], has been reported to up-regulate the expression of *CCHCR1* [21] and E2F1 [28, 29]. We observed prominent S phase arrest after a 24-hour treatment with 2 M CPT (Fig 2F; left), and importantly, both *CCHCR1* and *TCF19* were up-regulated (Fig 2F; right), suggesting that CPT-induced S-phase arrest could explain the up-regulation of both genes. This finding underscores the importance of understanding the cell cycle status when interpreting the effects of drugs and experimental conditions on gene expression.

## Discussion

The *CCHCR1* gene has received ample attention in clinical research in the last decade because of its potential association with psoriasis [30-32], alopecia areata [11], type-2 diabetes [12], and COVID-19 susceptibility [13]. Early genetic association studies showed controversial results for *CCHCR1* as the psoriasis susceptibility gene, possibly due to the small sample sizes, different populations and different statistical analyses being used [30-32]. As psoriasis is a complex multifactorial disease [33], it is also likely that one risk allele cannot fully explain the pathogenesis. In this study, we found that *CCHCR1* transcription is regulated by the G1/S-regulatory transcription factor E2F1, within a bidirectional promoter shared with *TCF19*. We hypothesized that this configuration allows synchronized induction of both genes during the G1/S transition of the cell cycle.

Human genome sequencing revealed an abundance (>10%) of divergently transcribed gene pairs whose transcription start sites are separated by less than 1000 bp [34]. A shared part of the intergenic fragment initiates and regulates transcription of the paired gene in both directions [34]. These genes are highly co-expressed [35] and may function in the same biological pathway such as DNA repair, cell cycle process, chromatin modification, metabolic process, and cell proliferation and differentiation [34, 36]. The cellular function of TCF19 is still vague. It is predicted as a transcription factor due to the presence of a putative trans-activating domain [37]. Mechanistically, TCF19 was shown to recognize H3K4me3 [38] to control the expression of glucose-responsive genes [38, 39]. Similar to CCHCR1, TCF19 has been associated with diseases, including diabetes [40-42], HBV-related chronic hepatitis B, cirrhosis, hepatocellular carcinoma [43], non-small cell lung cancer[44], squamous cell carcinoma of the head and neck (SCCHN) [45] and colorectal cancer [46]. It is currently not known whether CCHCR1 and TCF19 act on the same cellular pathway. One thing to note is both genes seem to be associated with diabetes. Interestingly, the head-to-head arrangement of *CCHCR1* and *TCF19* is also conserved in mouse, with a short intergenic region separated by only 291 bp. Further investigation into the genomic arrangement of *CCHCR1*-*TCF19* is essential for understanding the functional interplay between these genes, which may uncover shared pathways in disease processes and provide insights for targeted therapeutic strategies.

We and others have identified CCHCR1 as a novel component in cytoplasmic P-bodies and centrosomes [7, 8]. In this study, we demonstrated that *CCHCR1* is highly expressed during the G1/S transition and S phase of the cell cycle, potentially linking CCHCR1 activity to critical cell cycle events within these structures. P-bodies are phase-separated condensates in the cytoplasm responsible for RNA processing. The size and number of P-bodies dynamically change throughout the cell cycle, increasing in the S phase and disassembling upon mitotic entry [47]. Inside the P-bodies, CCHCR1 interacts with the mRNA decapping protein EDC4 [8]. Whether CCHCR1 participates in mRNA processing is still not known. In the centrosome, CCHCR1 controls cytoskeletal organization [7], and CCHCR1 knockdown causes centriole duplication defects and multipolar spindle formation [10]. Notably, centrosome duplication and centriole replication occur at the onset of the S phase [48], coinciding with the induction time of *CCHCR1* (Fig 2C). Further investigations into how CCHCR1 impacts the dynamics of both P-bodies and centrosomes throughout the cell cycle could provide vital insights into its cellular functions. In conclusion, our study unveils the transcriptional regulation of *CCHCR1* in the cell cycle through E2F1. Further exploration of its co-regulation with *TCF19* and its functions in P-bodies and centrosomes may unlock new avenues for therapeutic interventions in a range of medical conditions.

## Materials and Methods

### *In silico* analysis of the transcriptional regulation of *CCHCR1*

The intergenic region of *CCHCR1*-*TCF19* was analyzed by EMBOSS Newcpgreport [49] for the presence of CpG islands and by MatInspector [50] for putative transcription factor binding sites. Genes positively correlated with *CCHCR1* expression was identified using Genevestigator [26].

### Quantitative PCR (qPCR)

Total RNA from HeLa cells was isolated using TRIzol^®^ Reagent (Life Technologies, USA) according to the manufacturers instructions. Extracted RNAs were treated with DNase to remove genomic DNA using a TURBO DNA-free™ Kit (Life Technologies, USA), and were reverse transcribed to first-strand cDNAs with random primers using PrimeScript First Strand cDNA Synthesis Kit (Takara, Japan). Quantitative PCR (qPCR) was performed using Fast SYBR Green Master Mix (Life Technologies, USA) in a Step One Plus Real-time quantitative PCR system (Life Technologies, USA), with ROX as the reference dye. A final dissociation stage was performed after the thermocycles to verify the specificity of the PCR primers. All amplifications were done in triplicate. The relative expression of gene transcripts was normalized to *RPL13A* expression, and gene expression levels were calculated using the Pfaffl method [51]. The primer pairs used are listed in S4 Table.

### Overexpression of E2F proteins

To overexpress E2F1, 2, and 3, the coding sequences of the corresponding E2Fs were cloned from cDNA of HeLa cells into pEGFP-N1 vectors, which contain a strong CMV promoter. The plasmids were transiently transfected into HeLa cells using Fugene HD transfection reagent (Roche, Germany) for 48 hours to express the corresponding E2F-EGFP fusion proteins. Successful transfection was verified by the GFP expression of the cells.

### siRNA transfection

HeLa cells were seeded in a 6-well plate overnight before being transfected with 2.5 μM of siRNAs using Lipofectamine RNAiMAX transfection reagent (Invitrogen, USA) according to the manufacturers instructions. *CCHCR1, TCF19*, and *E2F1* mRNA were depleted using pre-designed Silencer^®^ Select siRNAs (Ambion, Life Technologies, USA). The negative control siRNA (siCTL) was Silencer^®^ Select Negative Control No. 1 siRNA (#4390843, Ambion, Life Technologies, USA). We observed no significant change of *CCHCR1* and *TCF19* mRNA level after 48 horus post-transfection of siCTL, while strong down-regulation of *CCHCR1* and *TCF19* mRNA was observed when treated with their specific siRNAs (S1 Fig).

### Cell cycle synchronization

Cells were synchronized at the G1/S transition by a double thymidine block. Briefly, HeLa cells at 25-30% confluency were incubated with 2 mM thymidine in complete DMEM medium for 18 hours, released into the thymidine-free complete medium for 9 hours, and incubated again with 2 mM thymidine in complete DMEM medium for another 17 hours. The cells were released from the block by washing with PBS and incubated with the complete DMEM medium.

### Flow cytometry

Cells were trypsinized, fixed in 70% ethanol, stained with propidium iodide, and analyzed by flow cytometry using LSRFortessa Cell Analyzer (BD Biosciences, USA).

### Dual luciferase assay

Promoter fragments containing the *CCHCR1*-*TCF19* intergenic region were cloned from HeLa cell genomic DNA and inserted into the firefly luciferase (Fluc) reporter vector pGL4.17 (Promega, USA), in either forward or reverse orientation. Firefly luciferase activity was measured to test the transcription activities of different *CCHCR1*-*TCF19* intergenic fragments at different phases of the cell cycle. In brief, HeLa cells (1×10^6^) were seeded in 24-well plates and incubated for 24 hours before transfection. Double transfections were performed using 200 ng total DNA containing 100 ng of Fluc reporter vector and 100 ng of Renilla luciferase (Rluc) vector (pRL-TKl; Promega). After 4 hours of transfection, cells were synchronized by double thymidine treatment and harvested at different time points after release from the block. Cells were lysed and luciferase activity was determined using a Dual-Luciferase Reporter Assay System (Promega) with a GloMax-96 Microplate Luminometer (Promega) as previously described [52]. To calculate the luciferase activity, the Fluc activity was first normalized to the Rluc activity in each individual experiment to correct for differences in transfection efficiency. Next, the Fluc/Rluc ratio of the empty vector control (pGL4.17) was subtracted from the Fluc/Rluc of each of the promoter fragments.

## Statistical analysis

All experiments were done in triplicate and data are presented as the mean value and standard deviation. Statistical analysis was performed with GraphPad Prism software version 7.0 (GraphPad Software, USA). A paired t-test was used to determine the significance of the data compared to the control. Statistical significance levels are denoted as: ^*\**^ P *≤* 0.05, ^*\*\**^ P *≤* 0.01 and ^*\*\*\**^ P *≤* 0.001; ns, not significant.

## Acknowledgments

We thank Prof. Z. X. Lin of the School of Chinese Medicine for technical assistance. This study was supported by the School of Life Sciences and School of Biomedical Sciences of CUHK.

## Supporting information

S1 Table. Putative transcription factor binding sites in the *CCHCR1*-*TCF19* bidirectional promoter.

S2 Table. Putative E2F binding motif in the *CCHCR1*-*TCF19* bidirectional promoter.

S3 Table. CCHCR1 co-expressed genes.

S4 Table. Primers for quantitative PCR.

S1 Fig. siRNAs control experiment.

