## Supporting information for "Cell cycle regulation of the psoriasis associated gene *CCHCR1* by transcription factor E2F1"

| Transcription factor | Matrix[A] | Detailed Matrix Information | Matrix similarity[B] | Strand | Position[C] |
|---|---|---|---|---|---|
| GABP | V$GABPB1.01 | GA repeat binding protein, beta 1 | 0.807 | (-) | -1208 to -1188 |
| | V$GABP.01 | GABP: GA binding protein | 0.889 | (+) | +79 to +99 |
| MYC | V$CMYC.02 | Myelocytomatosis oncogene (c-myc proto-oncogene) | 0.940 | (+) | -271 to -255 |
| NFY | V$NFY.04 | Nuclear factor Y (Y-box binding factor) | 0.925 | (+) | -913 to -899 |
| | V$NFY.04 | Nuclear factor Y (Y-box binding factor) | 0.939 | (+) | -710 to -696 |
| | V$NFY.04 | Nuclear factor Y (Y-box binding factor) | 0.962 | (-) | -257 to -243 |
| | V$NFY.03 | Nuclear factor Y (Y-box binding factor) | 0.864 | (-) | -184 to -170 |
| YY1 | V$YY1.02 | Yin and Yang 1 repressor sites | 0.944 | (-) | +299 to +321 |

**S1 Table. Putative transcription factor binding sites in the *CCHCR1-TCF19* bidirectional promoter.** [A]: MatInspector library: Matrix Family Library Version 9.2 (January 2015). [B]: A score of 1.00 indicates a perfect match to the matrix. A “good” match to the matrix has a score > 0.80. [C]: Numbers refer to the position related to the transcription start site (+1) of *CCHCR1* (NCBI Reference Sequence: NM_019052.3).

| Matrix[A] | Detailed Matrix Information | Matrix similarity[B] | Strand | Position[C] |
|---|---|---|---|---|
| V$E2F.02 | E2F, involved in cell cycle regulation, interacts with Rb p107 protein | 0.844 | (+) | -936 to -920 |
| V$E2F4.01 | E2F transcription factor 4, p107/p130-binding protein | 0.978 | (+) | -863 to -847 |
| V$E2F4_DP2.01 | E2F-4/DP-2 heterodimeric complex | 0.799 | (-) | -594 to -578 |
| V$E2F.03 | E2F, involved in cell cycle regulation, interacts with Rb p107 protein | 0.853 | (+) | -350 to -334 |
| V$E2F2.01 | E2F transcription factor 2 | 0.882 | (-) | -340 to -324 |
| V$E2F2.01 | E2F transcription factor 2 | 0.888 | (+) | -339 to -323 |
| V$E2F3.02 | E2F transcription factor 3 (secondary DNA binding preference) | 0.892 | (-) | -292 to -276 |
| V$RB_E2F1_DP1.01 | RB/E2F-1/DP-1 heterotrimeric complex | 0.766 | (+) | -289 to -273 |
| V$E2F4.01 | E2F transcription factor 4, p107/p130-binding protein | 0.967 | (-) | -279 to -263 |
| V$E2F4_DP1.01 | E2F-4/DP-1 heterodimeric complex | 0.974 | (-) | -233 to -217 |
| V$E2F4_DP1.01 | E2F-4/DP-1 heterodimeric complex | 0.965 | (+) | -232 to -216 |
| V$E2F3.02 | E2F transcription factor 3 (secondary DNA binding preference) | 0.863 | (+) | -230 to -214 |
| V$E2F1_DP2.01 | E2F-1/DP-2 heterodimeric complex | 0.861 | (-) | -205 to -189 |
| V$E2F.01 | E2F, involved in cell cycle regulation, interacts with Rb p107 protein | 0.858 | (-) | -168 to -152 |
| V$E2F.03 | E2F, involved in cell cycle regulation, interacts with Rb p107 protein | 0.887 | (+) | -133 to -117 |
| V$E2F3.01 | E2F transcription factor 3 | 0.903 | (+) | -30 to -14 |
| V$E2F3.01 | E2F transcription factor 3 | 0.864 | (-) | -29 to -13 |
| V$E2F.03 | E2F, involved in cell cycle regulation, interacts with Rb p107 protein | 0.870 | (+) | -13 to +4 |
| V$E2F4.01 | E2F transcription factor 4, p107/p130-binding protein | 0.979 | (+) | +6 to +22 |
| V$E2F4.01 | E2F transcription factor 4, p107/p130-binding protein | 1.000 | (+) | +99 to +115 |
| V$E2F4.01 | E2F transcription factor 4, p107/p130-binding protein | 0.969 | (+) | +480 to +496 |

**S2 Table. Putative E2F binding motif in the *CCHCR1-TCF19* bidirectional promoter.** [A]: MatInspector library: Matrix Family Library Version 9.2 (January 2015). [B]: A score of 1.00 indicates a perfect match to the matrix. A "good" match to the matrix has a score > 0.80. [C]: Numbers refer to the position related to the transcription start site (+1) of *CCHCR1* (NCBI Reference Sequence: NM_019052.3).

**Genevestigator microarray dataset: Perturbations**

| Rank | Gene | Description | Pearson's correlation coefficient | Up-regulated in G1/S or S phase? |
|---|---|---|---|---|
| 1 | *CCHCR1* | coiled-coil alpha-helical rod protein 1 | 0.6324975 | Yes (This study) |
| 2 | *CCHCR1* | coiled-coil alpha-helical rod protein 1 | 0.5560147 | Yes (This study) |
| 3 | *CCHCR1* | coiled-coil alpha-helical rod protein 1 | 0.5327725 | Yes (This study) |
| 4 | *RECQL4* | RecQ protein-like 4 | 0.44209898 | Yes [1] |
| 5 | *NRM* | nurim (nuclear envelope membrane protein) | 0.41889465 | NYD |
| 6 | *KIFC1* | kinesin family member C1 | 0.40744227 | NYD |
| 7 | *WRAP53* | WD repeat containing, antisense to TP53 | 0.39196926 | NYD |
| 8 | *ASF1B* | ASF1 anti-silencing function 1 homolog B (S. cerevisiae) | 0.39032698 | Yes [2] |
| 9 | *RAD54L* | RAD54-like (S. cerevisiae) | 0.3883079 | NYD |
| 10 | *EME1* | essential meiotic endonuclease 1 homolog 1 (S. pombe) | 0.38170868 | NYD |
| 11 | *ZDHHC12* | zinc finger, DHHC-type containing 12 | 0.38015762 | NYD |
| 12 | *PKMYT1* | protein kinase, membrane associated tyrosine/threonine 1 | 0.3774253 | Yes [3] |
| 13 | *TCF19* | transcription factor 19 | 0.37672827 | Yes [4] (This study) |
| 14 | *DDX11* | DEAD/H (Asp-Glu-Ala-Asp/His) box helicase 11 | 0.373824 | NYD |
| 15 | *RECQL4* | RecQ protein-like 4 | 0.37250307 | Yes [1] |
| 16 | *DDX11* | DEAD/H (Asp-Glu-Ala-Asp/His) box helicase 11 | 0.37225822 | NYD |
| 17 | *STRA13* | stimulated by retinoic acid 13 homolog (mouse) | 0.37041822 | Yes [5] |
| 18 | *TROAP* | trophinin associated protein (tastin) | 0.37041137 | NYD |
| 19 | *C16orf59* | chromosome 16 open reading frame 59 | 0.3693077 | NYD |
| 20 | *E2F1* | E2F transcription factor 1 | 0.36807078 | Yes [6] |
| 21 | *RDM1* | RAD52 motif 1 | 0.3665757 | NYD |
| 22 | *BRCA1* | breast cancer 1, early onset | 0.36540088 | Yes [7] |
| 23 | *MXD3* | MAX dimerization protein 3 | 0.36121294 | Yes [8] |
| 24 | *TONSL* | tonsoku-like, DNA repair protein | 0.3605641 | NYD |
| 25 | *FAM203A* | family with sequence similarity 203, member A | 0.35827172 | NYD |

**Genevestigator microarray dataset: Cell lines**

| Rank | Gene | Description | Pearson's correlation coefficient |
|---|---|---|---|
| 1 | *RNASEH2A* | ribonuclease H2 subunit A | 0.50694025 |
| 2 | *NELFE* | negative elongation factor complex member E | 0.49831852 |
| 3 | *DHX16* | DEAH-box helicase 16 | 0.49187893 |

| | | | |
|---|---|---|---|
| 4 | *KIF2C* | kinesin family member 2C | 0.4866221 |
| 5 | *BAG6* | BCL2 associated athanogene 6 | 0.4837167 |
| 6 | *TUBB* | tubulin beta class I | 0.47268584 |
| 7 | *CDC20* | cell division cycle 20 | 0.46996227 |
| 8 | *CHAF1B* | chromatin assembly factor 1 subunit B | 0.4682017 |
| 9 | *CDC25C* | cell division cycle 25C | 0.46737617 |
| 10 | *TCF19* | transcription factor 19 | 0.4619712 |
| 11 | *KIF4A* | kinesin family member 4A | 0.46129447 |
| 12 | *AURKB* | aurora kinase B | 0.45510256 |
| 13 | *PTTG1* | pituitary tumor-transforming 1 | 0.44862267 |
| 14 | *SPAG5* | sperm associated antigen 5 | 0.44852474 |
| 15 | *TUBBP2* | tubulin beta pseudogene 2 | 0.44831976 |
| 16 | *FKBPL* | FK506 binding protein like | 0.4475556 |
| 17 | *NRM* | nurim (nuclear envelope membrane protein) | 0.44644895 |
| 18 | *CSNK2B, XXbac-BPG32J3.22* | CSNK2B:casein kinase 2 beta | 0.44583622 |
| 19 | *TPX2* | TPX2, microtubule-associated | 0.441935 |
| 20 | *DONSON* | downstream neighbor of SON | 0.44076356 |
| 21 | *DSN1* | DSN1 homolog, MIS12 kinetochore complex component | 0.43877697 |
| 22 | *PARP2* | poly(ADP-ribose) polymerase 2 | 0.438576 |
| 23 | *ABCF1* | ATP binding cassette subfamily F member 1 | 0.43632245 |
| 24 | *GTF2H4* | general transcription factor IIH subunit 4 | 0.43618095 |
| 25 | *MEA1* | male-enhanced antigen 1 | 0.43477845 |
| **Genevestigator microarray dataset: Cancers** | | | |
| **Rank** | **Gene** | **Description** | **Pearson's correlation coefficient** |
| 1 | *TSEN54* | tRNA splicing endonuclease subunit 54 | 0.55687207 |
| 2 | *ASF1B* | anti-silencing function 1B histone chaperone | 0.5317818 |
| 3 | *POLD1, CTD-2545M3.6* | POLD1:polymerase (DNA directed), delta 1, catalytic subunit | 0.52976793 |
| 4 | *RECQL4* | RecQ like helicase 4 | 0.5265696 |
| 5 | *NRM* | nurim (nuclear envelope membrane protein) | 0.50892645 |
| 6 | *MCM2* | minichromosome maintenance complex component 2 | 0.50102276 |
| 7 | *ESPL1* | extra spindle pole bodies like 1, separase | 0.49512082 |
| 8 | *MYBL2* | v-myb avian myeloblastosis viral oncogene homolog-like 2 | 0.4910358 |
| 9 | *C19orf48, SNORD88B* | C19orf48:chromosome 19 open reading frame 48, SNORD88B:small nucleolar RNA, C/D box 88B | 0.4903525 |
| 10 | *CDCA8* | cell division cycle associated 8 | 0.4883732 |

| | | | |
|---|---|---|---|
| 11 | *POLE* | polymerase (DNA directed), epsilon, catalytic subunit | 0.48658773 |
| 12 | *TCF19* | transcription factor 19 | 0.4831117 |
| 13 | *CENPM* | centromere protein M | 0.46817404 |
| 14 | *ATAD3B* | ATPase family, AAA domain containing 3B | 0.46814716 |
| 15 | *RNASEH2A* | ribonuclease H2 subunit A | 0.4660578 |
| 16 | *POC1A* | POC1 centriolar protein A | 0.46444392 |
| 17 | *KIF2C* | kinesin family member 2C | 0.4628259 |
| 18 | *RNF26* | ring finger protein 26 | 0.45479703 |
| 19 | *RAD54L* | RAD54-like (S. cerevisiae) | 0.45234054 |
| 20 | *KIFC1* | kinesin family member C1 | 0.45149833 |
| 21 | *FANCA* | Fanconi anemia complementation group A | 0.44833374 |
| 22 | *RCCD1* | RCC1 domain containing 1 | 0.44649145 |
| 23 | *LAGE3P1* | L antigen family member 3 pseudogene 1 | 0.44513708 |
| 24 | *MYO19* | myosin XIX | 0.4434382 |
| 25 | *POLA2* | polymerase (DNA directed), alpha 2, accessory subunit | 0.44328704 |

**S3 Table. *CCHCR1* co-expressed genes.** Data retrieved using Genevestigator [9]. Top 25 positively correlated genes were shown. Repeated gene entry in the data is due to multiple microarray probes of the same gene being used in different studies. NYD: Not yet determined.

| Gene | Primer sequences (5' – 3') | Amplicon size (bp) |
|---|---|---|
| *RPL13A* | AGATGGCGGAGGTGCAG | 128 |
| | GTTGATGCCTTCACAGCGTA | |
| *CCHCR1* | GGAAGAACTTGGAAGAGGGG | 141 |
| | GAGACTTCTCCAAGCCCTCA | |
| *TCF19* | GGTGATGACTGGAGGGTCAG | 111 |
| | CAGGAGGTCTCCATCACTCA | |
| *E2F1* | CCAGGAAAAGGTGTGAAATC | 74 |
| | AAGCGCTTGGTGGTCAGATT | |

**S4 Table. Primers for quantitative PCR.**

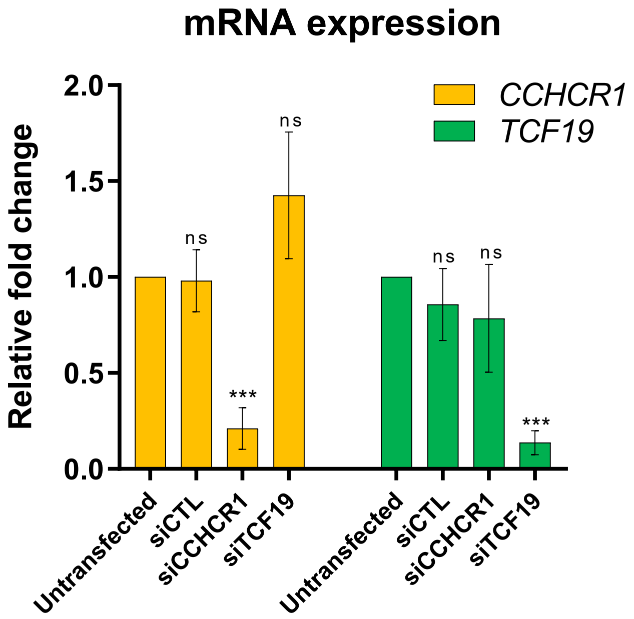

**S1 Fig. siRNAs control experiment.**
